# A new look at multi-stage models of cancer incidence

**DOI:** 10.1101/243972

**Authors:** Tyler Lian, Rick Durrett

## Abstract

Multi-stage models have long been used in combination with SEER data to make inferences about the mechanisms underlying cancer initiation. The main method for studying these models mathematically has been the computation of generating functions by solving hyperbolic partial differential equations. Here, we analyze these models using a probabilistic approach similar to the one Durrett and Moseley [7] used to study branching process models of cancer. This more intuitive approach leads to simpler formulas and new insights into the behavior of these models. Unfortunately, the examples we consider suggest that fitting multi-stage models has very little power to make inferences about the number of stages unless parameters are constrained to take on realistic values.

## 1 Introduction

Investigation of the age distribution of cancer incidence goes back to the middle of the 20th century. Fisher and Holloman [10] and Nordling [20] found that within the 25–74 age range, the logarithm of the cancer death rate increased in direct proportion to the logarithm of the age, with a slope of about six on a log-log plot. Nordling grouped all types of cancer together and considered only men, but the pattern persisted when Armitage and Doll [2] separated cancers by their type and considered men and women separately.

Nordling [20] suggested that the slope of six on a log-log plot would be explained if a cancer cell was the end result of seven successive mutations. There was no model underlying that conclusion, just the observation that if one sums *k* exponential random variables with rate *μ_i_*, i.e., with probability density 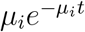, then when *t* is much smaller than the mean of the sum 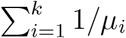, then “the probability that the *k*th change occurs in the short time interval (*t, t + dt*) is asymptotically

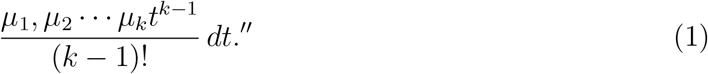

Later Armitage [1] gave a rigorous proof of this result.

A few years later, Armitage and Doll [3] wrote that the hypothesis described in the previous paragraph was “however, unsatisfactory in that there was no direct experimental evidence to suggest that carcinogenesis was likely to involve more than two stages.” Because of this, they introduced in [3] a two-stage model in which ordinary cells (type 0) mutate at rate *μ*_1_ into initiated (type 1) cells that grow at exponential rate λ and mutate at rate *μ*_2_ into malignant cells (type 2). The two-stage model has been thoroughly analyzed in the literature, see e.g. [12], [13]

In the studies cited above, the stages were unspecified events. That changed in 1971 with Knudson’s study of retinoblastoma [14]. Based on observations of 48 cases of retinoblastoma and published reports, he hypothesized that the disease is a cancer caused by two mutational events. In the dominantly inherited form, one mutation is inherited via the germinal cells. In the nonhereditary form both mutations occur in somatic cells. The underlying gene, named RB1, was found 15 years later. In currentl terminology, RB1 is a *tumor suppressor gene*. Trouble begins when both copies are knocked out.

Colorectal tumors provide an excellent system in which to search for and study the genetic alterations involved in the development of cancer because tumors of various stages of development, from very small adenomas to very large carcinomas, can be obtained for study. The initiating event is thought to involve the inactivation of the tumor suppressor gene APC (adenomatous polyposis coli). As in retinoblastoma, an inherited germ line mutation in this gene causes greater risk of disease. Individuals with this mutation have numerous polyps form early in their lives, mainly in the epithelium of the large intestine.

In 1990, Fearon and Vogelstein [8] found a second piece of the puzzle when they noted that approximately 50% of colorectal carcinomas, and a similar percentage of adenomas greater than 1 cm have mutations in the RAS gene family, while only 10% of adenomas smaller than 1 cm have these mutations. In the modern terminology, the members of the RAS family are *oncogenes*. A mutation to a single allele is sufficient for progression. The analysis in [8] also suggested a role for TP53 (which produces the tumor protein p53) in the progression to cancer. The protein p53 has been described as ”the guardian of the genome” because of its role in conserving stability by preventing genome mutations. TP53 has since been implicated in many cancers, see [11] and [25].

Combining the ideas in the last two paragraphs leads to a four (or five) stage description for colon cancer that is described for example in the books of Vogelstein and Kinzel [24], and Frank [9]. In 2002, Leubeck and Moolgavar [16] developed a mathematical model in order to fit the age-incidence of colorectal cancer. We will describe the model in detail in the next section. They tried models with *k* = 2, 3, 4, 5 stages and found that the four-stage model gave the best fit. The techniques developed in [16] have been applied to study a number of other cancers. See e.g. [17], [18], [19].

## 2 Analytic approach

In the *k*-stage model there is a fixed number of stem cells, *N*, each of which mutates at rate *μ*_0_ to become a type 1 cell, so cells of type 1 are born at times of a Poisson process with rate *γ* = *Nμ*_0_. Cells of types *i* = 1,… *k* − 2 are pre-initiated cells that mutate at rate *μ_i,k_* to become a cell of type *i* + 1. Cells of type *k −* 1 are initiated cells that divide into two at rate *α*, die at rate *β*, where λ = *α − β* > 0, and mutate at rate *μ_k_*_−1,_*_k_* to become malignant (type *k*). Let 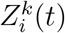 be the number of type *i* cells at time *t* in the *k*-stage model. Let 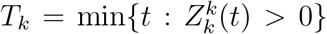 be the time of appearance of the first malignant cell in the k-stage model. Here we will be interested primarily in

- the survival function *H_k_*(*t*) = *P*(*T_k_ > t*), which gives the fraction of individuals that are cancer free at time *t*
- hazard rate 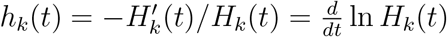, which gives the rate at which healthy individuals become sick at time *t*.

The traditional approach to studying the *k*-stage model, as explained for example in the supplementary materials of [16], has been to let

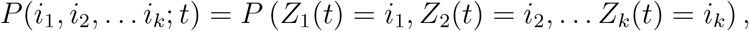

define the generating function

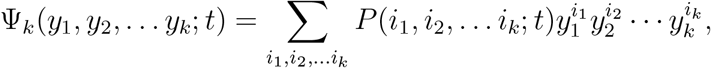

and compute Ψ*_k_* by solving a hyperbolic PDE using the method of characteristics.

To explain this approach, we will consider the case *k* = 2 and write

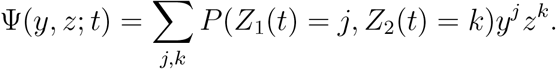

Transition rates are given in the following table.

| at rate | transition | g.f. change |
| --- | --- | --- |
| $\gamma$ | $j \rightarrow j + 1$ | $\Psi \rightarrow y_1 \Psi$ |
| $\alpha j$ | $j \rightarrow j + 1$ | $\Psi \rightarrow y_1 \Psi$ |
| $\beta j$ | $j \rightarrow j - 1$ | $\Psi \rightarrow \Psi / y_1$ |
| $\mu_{1,2} j$ | $k \rightarrow k + 1$ | $\Psi \rightarrow y_2 \Psi$ |

From the table we get

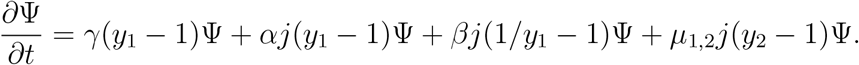

Using the identity *j*Ψ = *y*_1_∂Ψ/∂*y*_1_ this becomes

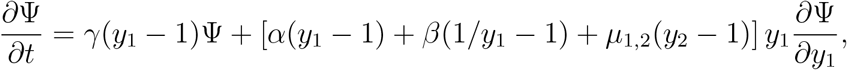

which rearranges to become

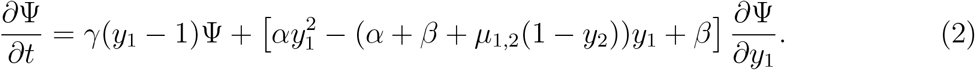

To find the generating function Ψ*_k_*(*z*_1_, *z*_2_,… *z_k_* : *t*), one uses the fact that the solution is constant along characteristic curves, i.e.,

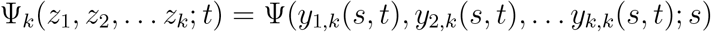

where the *y_i,k_*(*s, t*) satisfy the characteristic equations

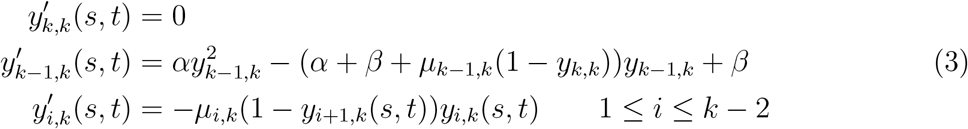

and the derivative is taken with respect to the *s* variable. The generating function can then be found from

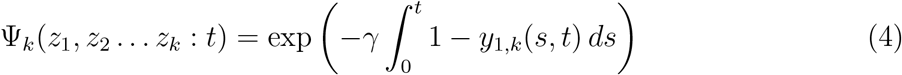

and the survival function 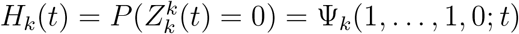.

### 2.1 Solving the equations

To begin to solve the equations in (3), we note that 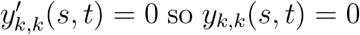 is constant. We write *S*_1,_*_k_*(*t*) = *y_k_*_−1,_*_k_*(*t*) since it is the first step in solving the system of equations. The subscript *k* is needed because the mutation rates that enter into the differential equations (3) depend on the number of stages. Throughout this paper we when we write *S_i,k_*(*t*) it is assumed that *z_k_* = 0 and *z_i_* = 1 for 1 ≤ *i* < *k*. Changing notation we want to solve

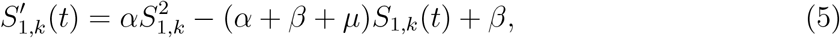

where *μ* = *μ_k−_*_1_*_,k_* and *S*_1_*_,k_*(0) = 1. The quadratic equation *αx*^2^ − (*α* + *β* + *μ*)*x* + *β* = 0 has two roots *q* > 1 > *r* > 0. See (31). Solving (5), see (34), gives

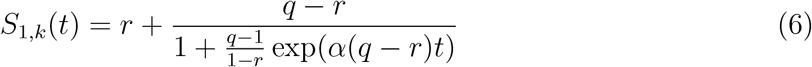

Having solved for *S*_1,_*_k_* (*t*) the other *S_i,k_* (*t*) = *y_k−i,k_*(*t*) can be found by induction:

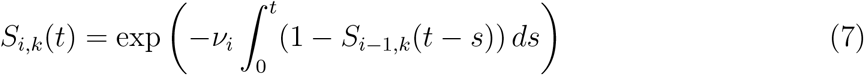

where in the *k*-stage model *v_i_* = *μ_k−i,k_*. The computation of the *S_i,k_*(*t*) is not easily found in the literature, so we will give the details in Section 7.

While the recursion in (7) was derived by the method of characteristics, it has a simple probabilistic interpretation. Each individual of type *k* − *i* gives birth to individuals of type *k* − *i* + 1 at times of a rate *v_i_* Poisson process. A type *k* − *i* + 1 born at time s will give rise to a malignant cell with probability 1 − *S_i_*_−1_(*t* − *s*). The number of type *k* − *i* + 1 individuals that are successful in doing this has a Poisson distribution with mean

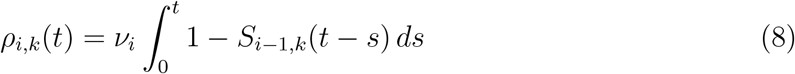

so the probability none of the type *k* − *i* + 1 individuals are successful in creating a malignant cell is *S_i,k_*(*t*) = exp(−*ρ_i,k_*(*t*)).

### 2.2 Hazard rate formulas

It is clear from (4) that *H_k_* = *S_k,k_* so we have

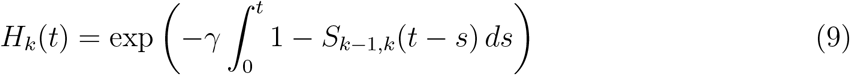

Using (9) and changing variables *r* = *t* − *s* before differentiating it follows that

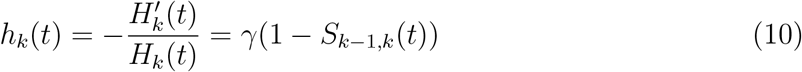

so we do not have to evaluate *H_k_*(*t*) to find *h_k_*(*t*).

Using (10) with (35) gives

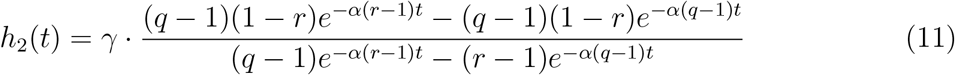

Using (37) we have

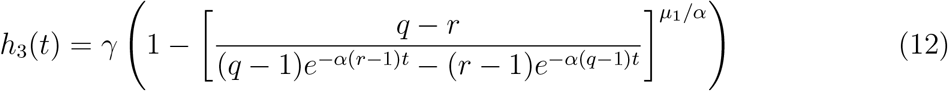

When it comes to the fourth stage, the possibility to compute the integral in (7) breaks down and (39) gives

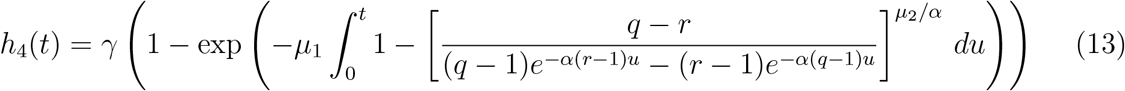

## 3 Probabilistic approach

We begin by giving probabilist interpretations for some of the computations above. To explain the differential equation for *y_k−_*_1_*_,k_*, note that if we ignore mutation then each individual of type *k* − 1 initiates a linear birth and death process *L*(*t*) in which the number of individuals increases from *m* → *m* + 1 at rate αm and decreases from *m* → *m* − 1 at rate *βm*, where *α* > *β*.

### Theorem 1.

*As t* → ∞, *e*^−λt^*L*(*t*) → *W with P*(*W* = 0) = *β*/*α* = *P*(*L*(*t*) = 0 *for some t* > 0) *and*

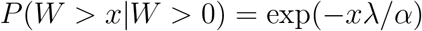

*i. e*., *if we condition on non-extinction then W has an exponential density with rate* λ/*α*. *If we let V*_0_ = (*W*|*W* > 0) *then*

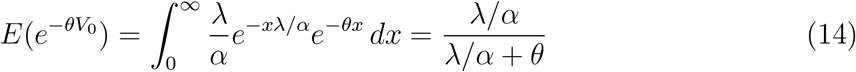

### Proof.

We will sketch the proof since it contains details that will be useful later. For more details see Section 3 of [6]. It is well know that if we start with *L*(0) = 1 then the generating function *F*(*x, t*) = *Ex^L^*^(^*^t^*^)^ satisfies

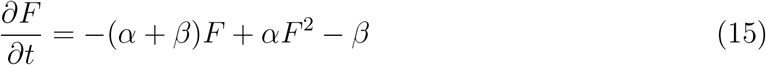

with boundary condition *F*(*x*, 0) = *x*. This equation can be solved with the result that

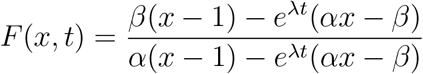

where λ = *α* − *β* is the exponential growth rate. By considering what happens on the first step (which is a birth with probability *α*/(*α* + *β*) and a death with probability *β*/(*α* + *β*)) we can conclude that the probability *ρ* that the process dies out satisfies

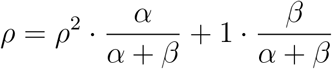

The extinction probability is the root which is < 1, i.e., *ρ* = *β*/*α*.

Comparing with (15) we see that (5) has an additional term −*μy*(*t*). In probabilistic terms, this corresponds to killing the process at rate *μm* when there are *m* individuals in the branching process. Let 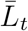 be the birth and death chain conditioned not to die out. Using this observation and Theorem 1, the probability of no malignant cell by time *t* in 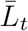 is

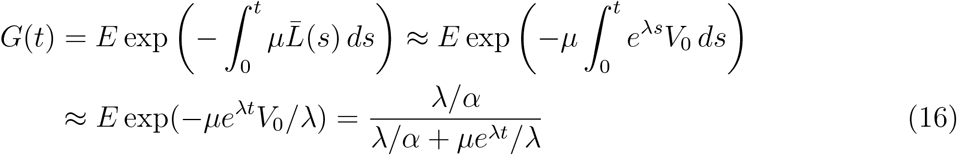

where in the last step we have used (14). When the branching process dies out it does that quickly so the probability of a mutation is small. From this it follows that

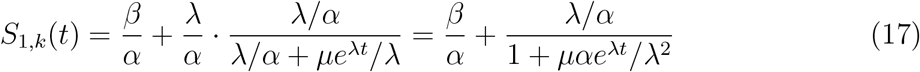

To compare with (6) we need to change notation. Equations (41) and (42) imply that

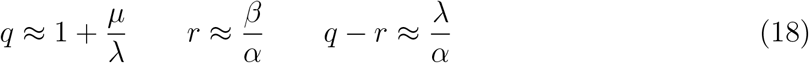

Using these in (6) we have

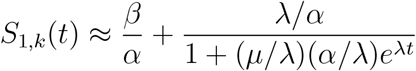

which agrees with the new formula in 17.

To compute the survival function *H_k_*(*t*) from this we start at 1 and work up to type *k* − 1. Let *η_j_*(*s*) be the rate at which type *j*’s born at time *s*. Integrating we find

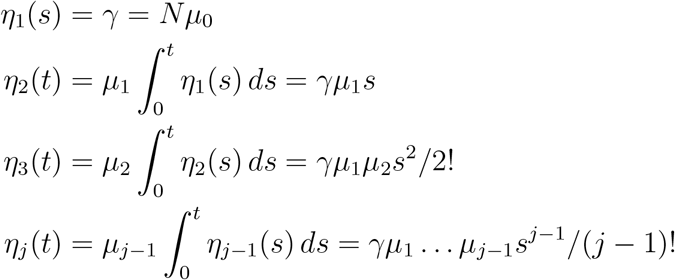

We call a type *k* − 1 family that does not die out “successful.” Using Theorem 1, the probability that a type *k* − 1 is successful is λ/*α*. On the event that a type *k* − 1 is produced in [0, *T*], the time it is born will be distributed as *η_k−_*_1_(*s*)λ/*α*. Recalling that 1 − *G*(*t* − *s*) is the probability a successful birth and death process conditioned to not die out gives rise to a malignant cell and using the reasoning that led to (8), the times of successes will be roughly a Poisson process so

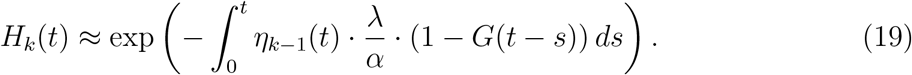

Using this approach we find (see Section 8 for details) that

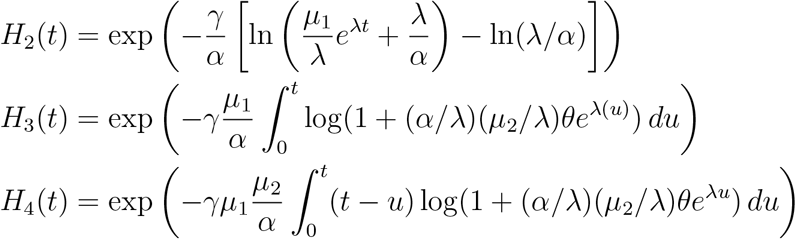

Differentiating with respect to *t*, and using 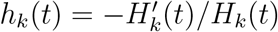

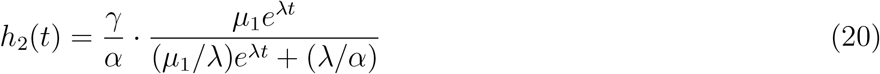

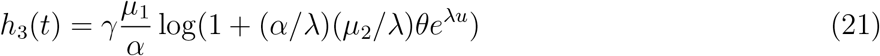

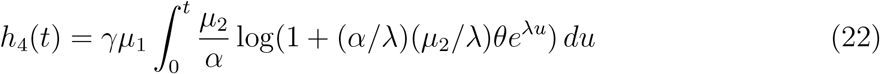

At first glance the new formulas look much different than the ones from the analytic approach. However, a closer look shows they are closely related. When *k* = 2, *η_k_*_−1_ = γ and (17) implies

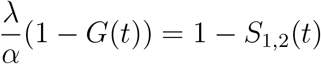

so the formulas for *h*_2_(*t*) agree. Comparing the two formulas for *h*_3_(*t*) and *h*_4_(*t*) we see that it is enough to argue

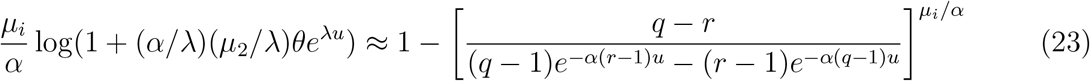

## 4 The two-stage model

The formula for the hazard rate in the two stage case, given in (11), is

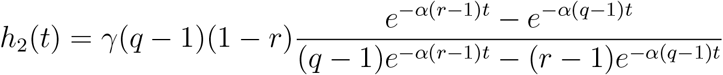

If we introduce *P* = *γ*(*q* − 1) and *Q* = *α*(*q* − 1) we can rewrite it as

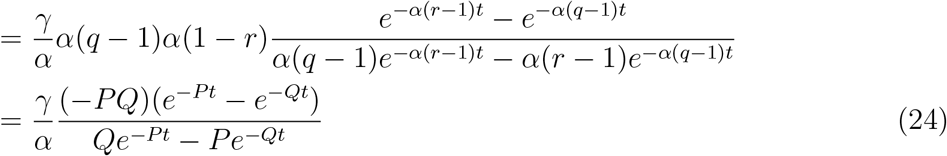

Note that the two-stage model has five parameters γ, *μ*_1_, *μ*_2_, *α*, and *β* but the new formula for the hazard rate has only three *γ*/*α*, *P* = *α*(*r* − 1), and *Q* = *α*(*q* − 1). In the terminology of statistics we have an identifiability problem, i.e., not all the parameters in the model can be estimated.

**Table 1:** Fits of the two stage model hazard rate given in (24) to thyroid cancer in white females by Meza and Chang [18] and to peritoneal mesothelioma by Moolgavkar, Meza, and Turim [17].

| parameter | [18] | [17] |
| --- | --- | --- |
| $\gamma/\alpha$ | $3.87 \times 10^{-4}$ | $3.17 \times 10^{-5}$ |
| $-P$ | 0.259 | 0.11 |
| $Q$ | $8.22 \times 10^{-4}$ | $1.78 \times 10^{-4}$ |

Figure 1 gives a picture of *h*_2_(*t*) for the parameters of [18]. It should be obvious from the picture that as *t* → ∞, *h*_2_(*t*) converges to a limit. Since *P* < 0 < *Q* letting *t* → ∞ in (24)

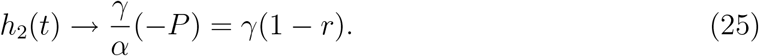

Figure 2 shows the fit of the two stage model to peritoneal mesotheliomas in SEER data from 1973–2005. Parameters are given in Table 1. This time the asymptote has not been reached by age 85. The dotted line gives the fit of the Armitage-Doll formula (1) *Ct^k^* to the data with *C* = 1.75 × 10^−11^ and *k* = 2.79. Visually the second fit is worse. This is confirmed by the values of Akaike Information Criterion scores. Interested readers can consult [17] for further details.

**Figure 1:**
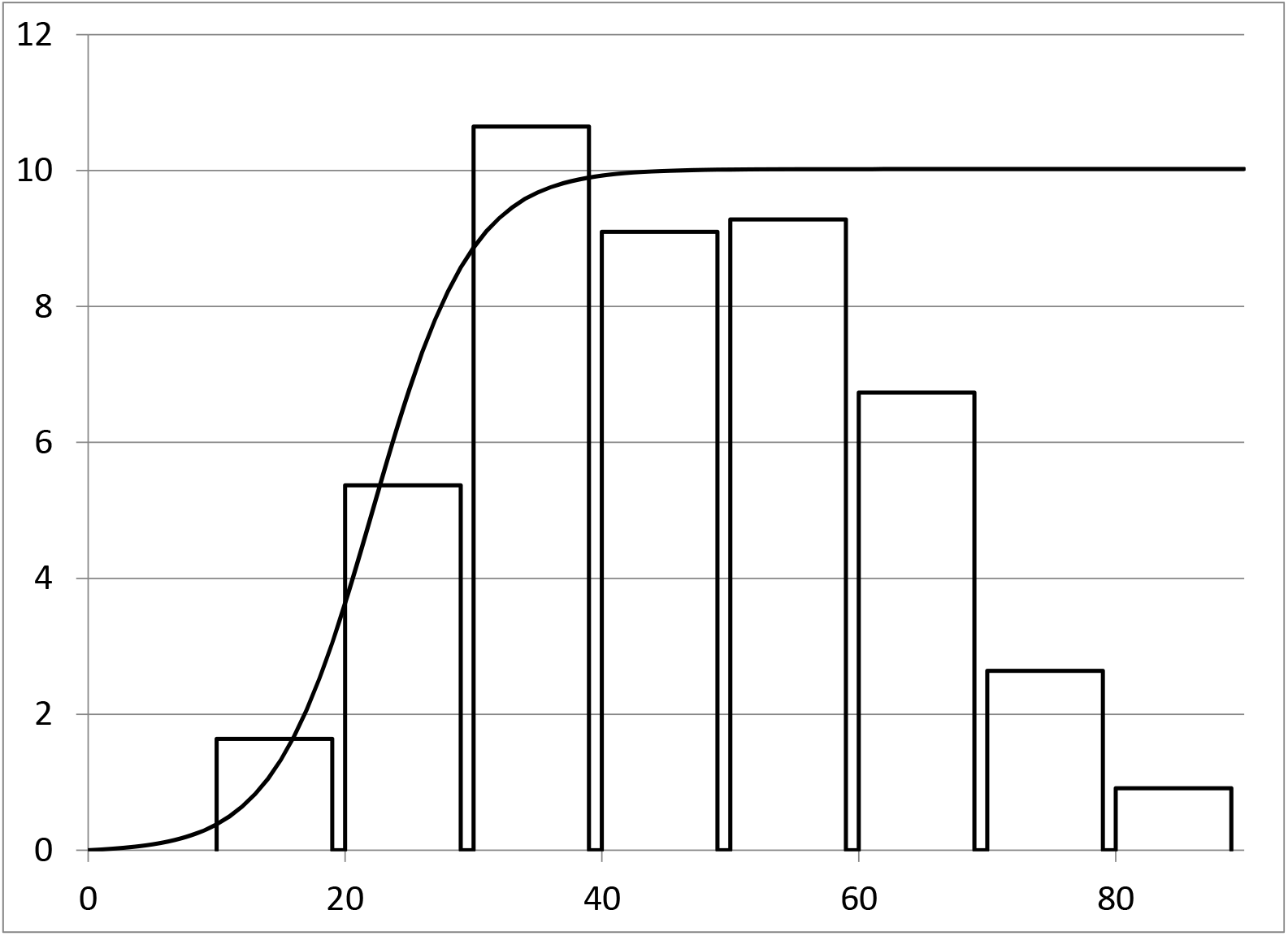
A graph of *h*_2_(*t*) for the thyroid cancer parameters. *x* axis is age in years. *y* axis is cases per 100,000 per year. The asymptotic value, which is 10.02 by (25) is reached at age ≈ 40. For comparison, we give a histogram of age at diagnosis in 508 individuals in a TCGA study [22]. If one transforms the data so that it is cases per 100,000 individuals in each age group, the model fits the data. See Figure 4 in [18].

**Figure 2:**
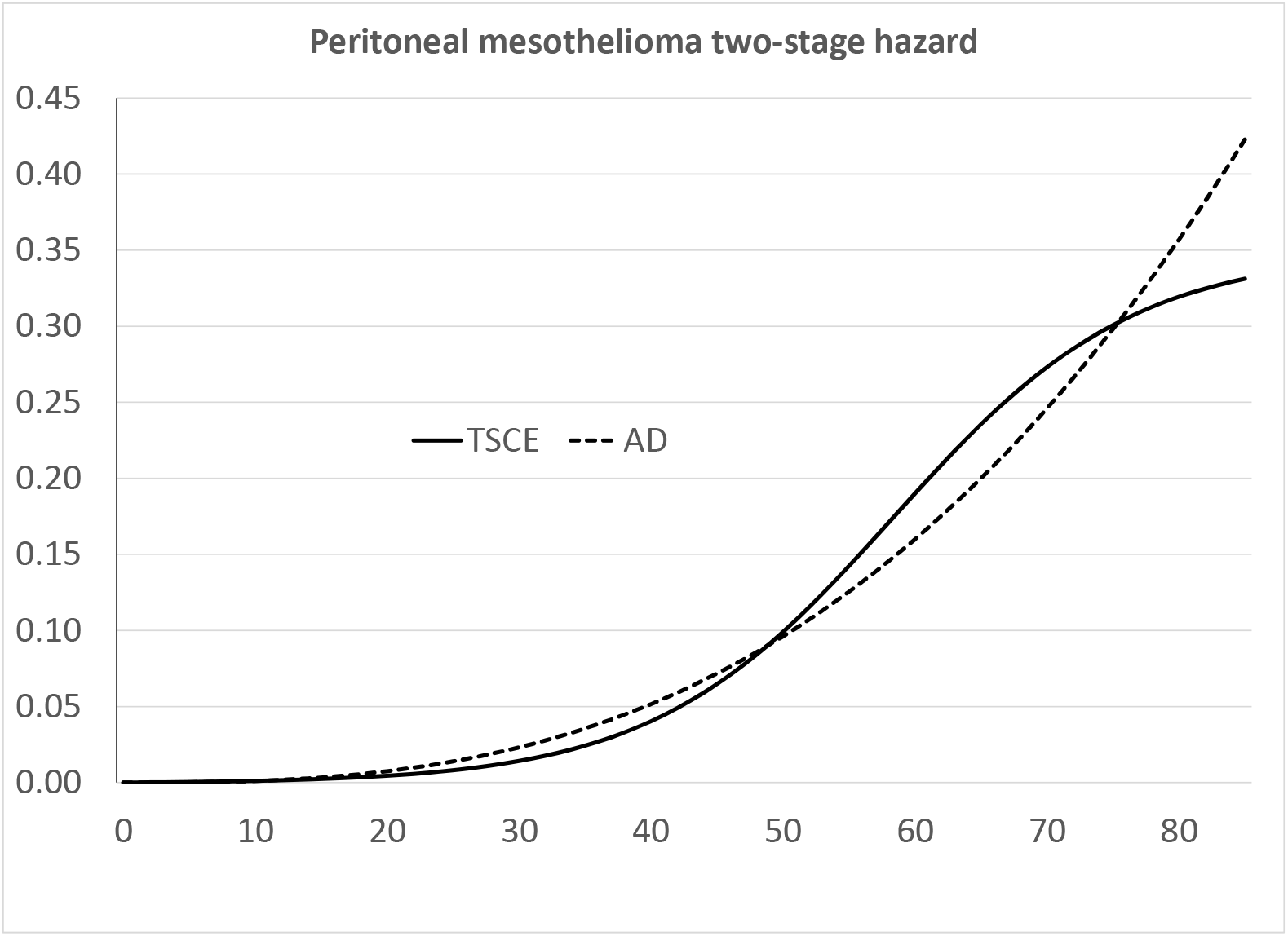
The solid line is a graph of *h*_2_(*t*) for the peritoneal mesothelioma parameters. The dotted line is a fit of the Armitage-Doll model with *k* = 2.79. *x* axis is age in years. *y* axis is cases per 100,000 per year. The asymptotic value has not been reached by age 85.

## 5 The three-stage model

Using (12) and changing variables *P* = *α*(*r* − 1) and *Q* = *α*(*q* − 1) it follows that the hazard rate is

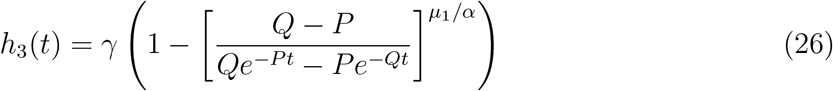

Meza at al [19] show that *h*_3_ is asymptotically linear. To state their results we need two definitions. The probability that the birth and death processes does not die out is

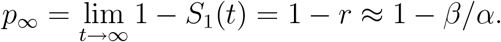

by (6) and (18). Let *T*_2,3_ be the time to malignancy of a single type 2 clone in the 3-stage model conditional on it not becoming extinct.

### Theorem 2.

*If t is large and t* ≪ 1/*μ*_1_*p*_∞_ *then*

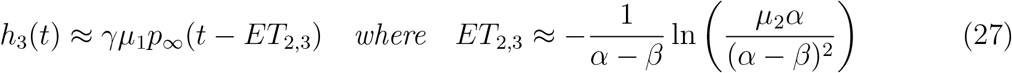

To better understand the formula for *h*_3_(*t*), and to check the accuracy of the linear approximation, it is useful to have concrete examples.

**Table 2:** Meza et al [19] estimated parameters for the 3-stage model and 4-stage model (described in the next section) for pancreatic cancer in men.

| name | parameter | 3-stage fit | 4-stage fit |
| --- | --- | --- | --- |
| slope | $\gamma \mu_1 p_\infty$ | $3.9 \times 10^{-5}$ | $4.68 \times 10^{-5}$ |
| $-P$ | $\alpha - \beta$ | 0.179 | 0.192 |
| | $\mu_2/\alpha$ | NA | 0.401 |
| $Q$ | $\alpha \mu_{k-1}/(\alpha - \beta)$ | $1.38 \times 10^{-5}$ | $2.06 \times 10^{-5}$ |
| | | $ET_{2,3} = 52.9$ | $ET_{2,4} = 57.9$ |

To be able to compute the hazard function we need a value for *α*. Meza at al [19] suggest *α* = 9 cell divisions per year and say that the fit is not sensitive to the value of α chosen. In pancreatic cancer, when *α* = 9,

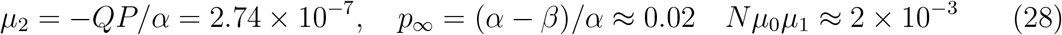

If we take *μ*_0_ = *μ*_1_ = 10^−6^ then *N* = 2 × 10^9^, and *μ*_1_/α =1.1 × 10^−7^. Figure 3 gives a graph of *h*_3_(*t*) (and *h*_4_(*t*)) for the pancreatic cancer parameters. 1/*μ*_1_*p*_∞_ = 5 × 10^9^ years so the condition *t* ≪ 1/*μ*_1_*p*_∞_ holds. As the graph shows the straight line approximation is good for *t* ≥ 65.

**Figure 3:**
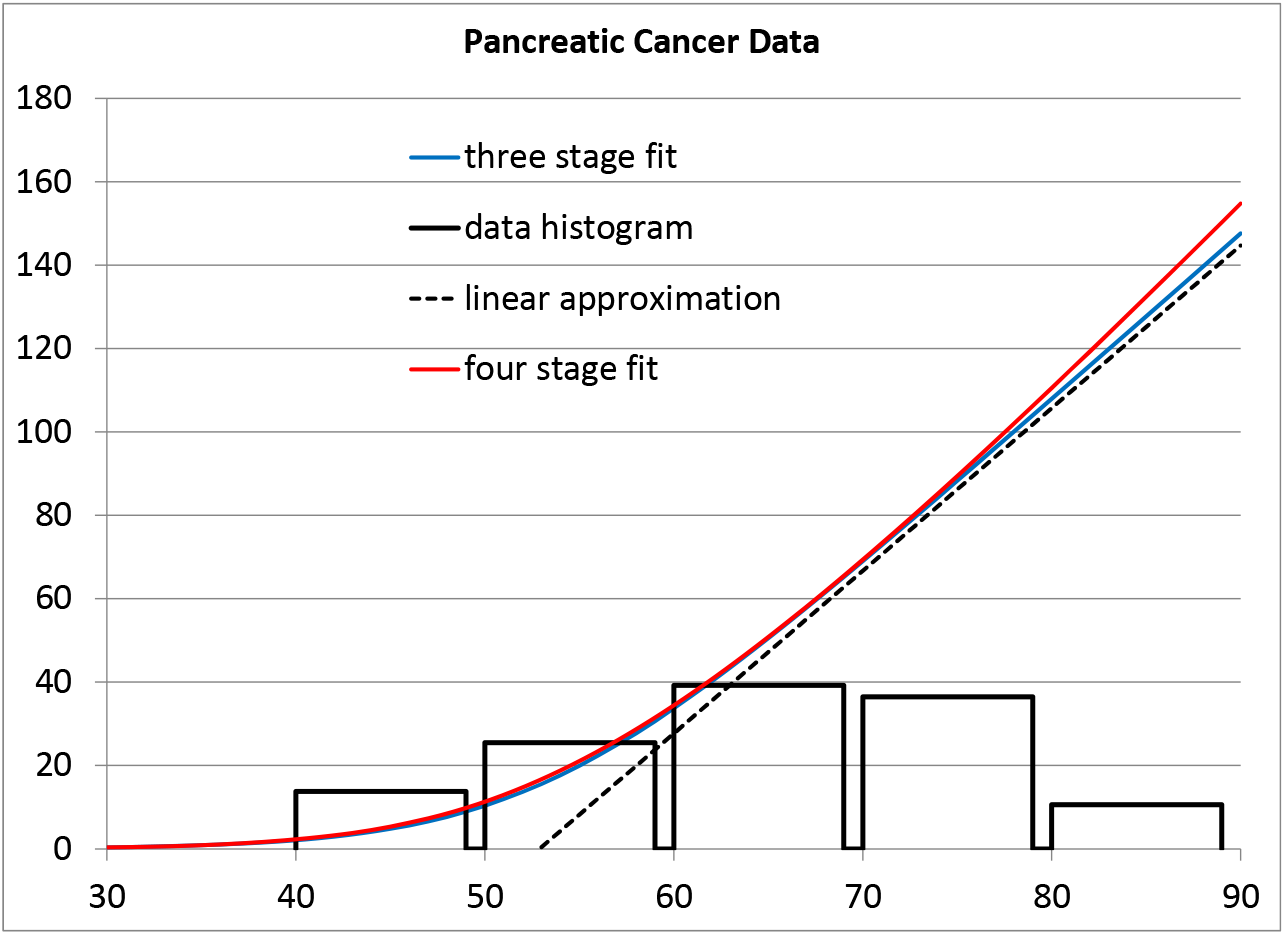
Graphs of *h*_3_(*t*) and *h*_4_(*t*) for the pancreatic cancer parameters. *x* axis is age in years. *y* axis is cases per 100,000 per year. Straight line is the linear approximation to the three stage model (27). The bar graph gives the age at diagnosis for 186 patients in the TCGA study of pancreatic cancer [4]. Again if one transforms the data to be cases per 100,000 in each age group, the theoretical curve fits the data, see Figure 5 in [19].

## 6 The four-stage model

### 6.1 Hazard rate

Using (13) and changing variables *P* = *α*(*r* − 1) and *Q* = *α*(*q* − 1)

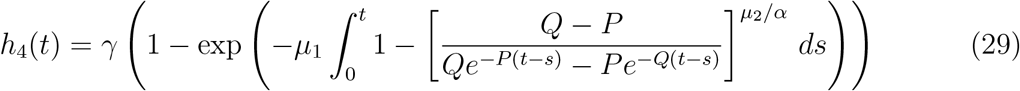

Let *T*_2,4_ be the time for a single type 2 clone to produce a malignant cell in the four-stage model and let

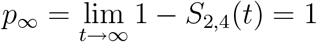

since a type 2 will give rise to infinitely many type 3’s, and one of these will start a branching process that does not die out.

The asymptotic behavior of the hazard rate is constant in the two-stage case and linear in the three-stage case. One might naively guess that in the four-stage case it is asymptotically quadratic, but the simple proof given below shows it is asymptotically linear. It should be clear from the proof that this holds for any *k* ≥ 3.

#### Theorem 3.

*When t is large and t* ≪ 1/*μ*_1_

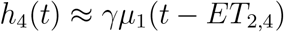

##### Proof of Theorem 3.

When *μ*_1_*t* is small

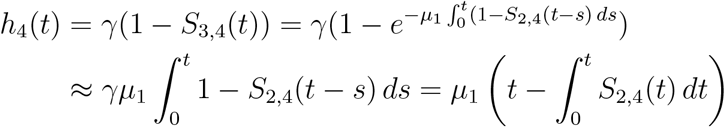

Using a well-known formula for expected value

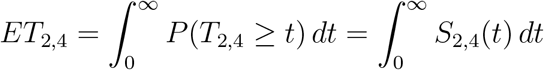

Combining the last two equations gives the desired result.

Note: due to the complexity of the formula for *S*_2,4_(*t*) given in (36), we do not have a formula for *ET*_2,4_. However, it is easy to compute numerically.

Leubeck and Moolgavkar [16] estimated parameters for the 2, 3, 4, and 5 stage models for colorectal cancer in women. As Figure 4 shows the fits from the four models are all very good. To explain how this could happen, we take a look at the parameters used in fitting. *N* = 10^8^, *α* = 9. Note that in the four stage model, *μ*_2_ = 6.3, and in the five-stage model *μ*_3_ = *μ*_4_ = 0.9. These large values speed these processes up, effectively eliminating one and two stages respectively. In the other direction in the two stage model the very slow mutation rates *μ*_0_ = 4.5 × 10^−9^ and *μ*_1_ = 1.44 × 10^−7^ effectively add a stage. Thus if we judge the fitted model by the size of the mutation rates, it seems that the three-stage model gives the best fit.

**Figure 4:**
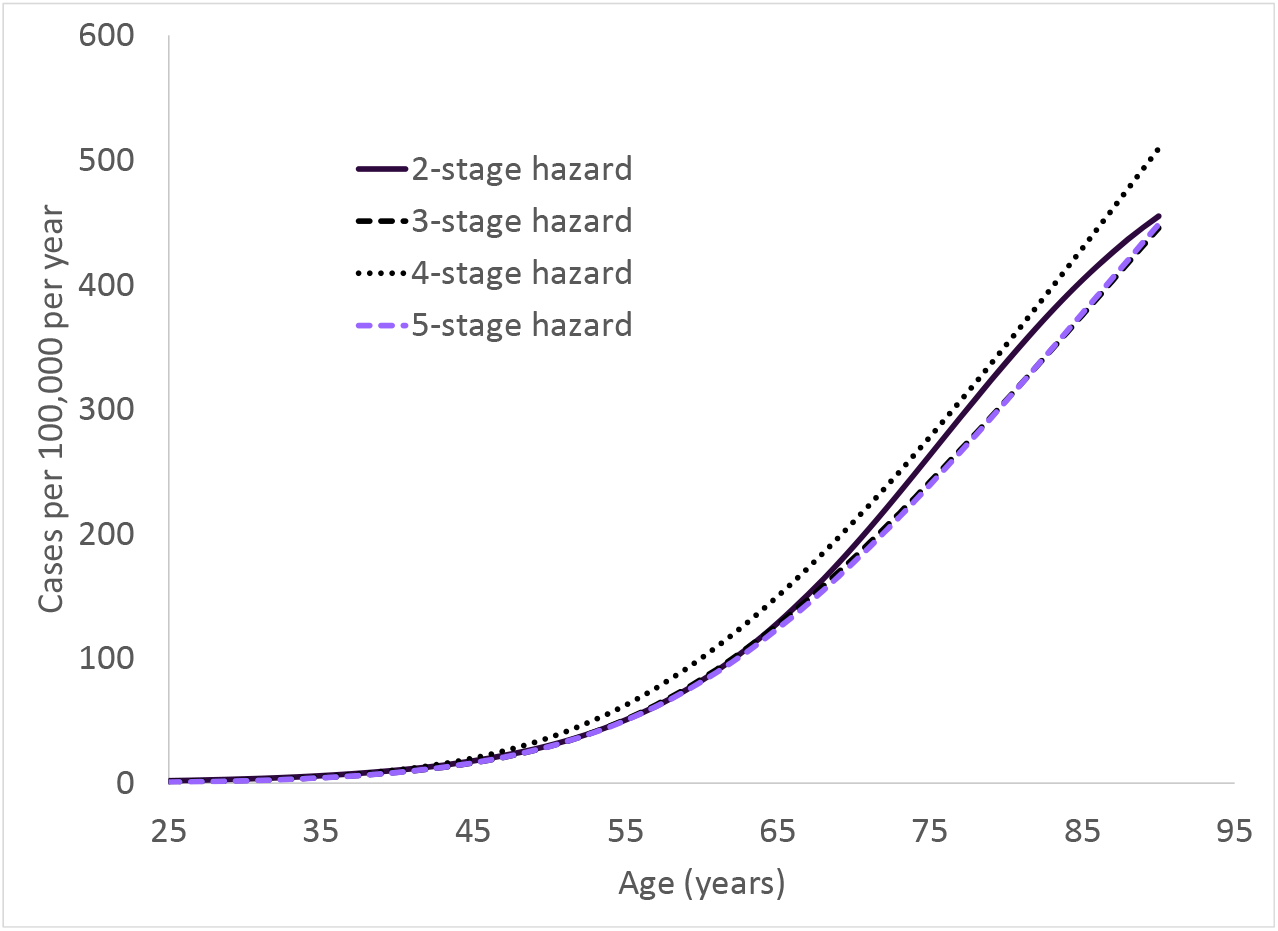
A comparison of the fitted values of the hazard functions for the two, three, four, and five stage models of [16]. The three and five stage fits are almost identical so you can only see three curves on the graph.

**Table 3:** Parameter values in four fits of colon cancer data from [16].

|  | 2-stage | 3-stage | 4-stage | 5-stage |
| --- | --- | --- | --- | --- |
| $\mu_0$ | $4.5 \times 10^{-9}$ | $1 \times 10^{-5}$ | $1.3 \times 10^{-6}$ | $1.3 \times 10^{-6}$ |
| $\mu_1$ | $1.44 \times 10^{-7}$ | $1 \times 10^{-5}$ | $1 \times 10^{-6}$ | $1 \times 10^{-6}$ |
| $\mu_2$ | — | $8.77 \times 10^{-7}$ | 6.3 | 0.9 |
| $\mu_3$ | — | — | $1.333 \times 10^{-6}$ | 0.9 |
| $\mu_4$ | — | — | — | $1.89 \times 10^{-6}$ |
| $P$ | -0.11 | -0.13 | -0.13 | -0.11 |
| $Q$ | $2.64 \times 10^{-5}$ | $6.08 \times 10^{-5}$ | $9.23 \times 10^{-3}$ | $1.545 \times 10^{-4}$ |

It is interesting to note that Tomasetti et al [23] have arrived at the conclusion colon cancer is a three-stage process by a completely different reasoning. They compared patients with and without a mismatch repair deficiency. They found that the latter group has 7.7 to 8.8 times as many mutations, versus a 114.2 fold increase in colon cancer rates, and argued that the increase would be more substantial if the process had four-stages. See pages 119–120 in [23] for more details and an analysis of lung adenocarcinomas.

## 7 Computing *S_i,k_* (*t*): analytic approach

Let *q* > 1 > *r* be the roots of the quadratic equation

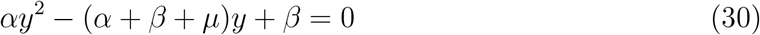

that is,

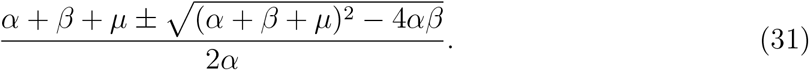

If we write *y*(*t*) = *S*_1,_*_k_*(*t*) then the differential equation (5) can be written as

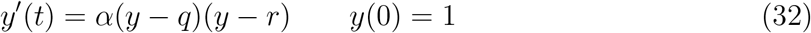

From this we see that *y*(*t*) is decreasing and will converge to r as *t* → ∞. Rearranging (32), we have

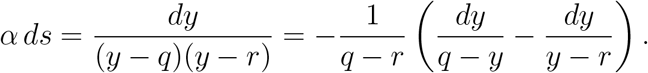

Here we have written the right-hand side to avoid taking the logarithm of a negative number in the next step. Multiplying both sides by *q* – *r* and then integrating from 0 to *t*, we have for some constant *D*

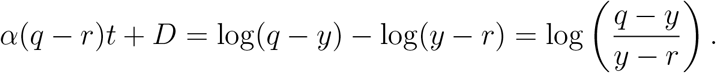

Exponentiating we have

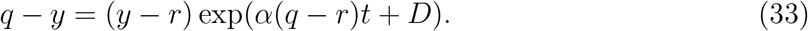

Solving for *y* gives

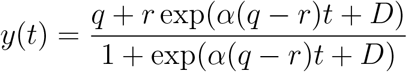

Using (33) and recalling *y*(*t*) = 1 we have *e^D^* = (*q* − 1)/(1 − *r*) which implies

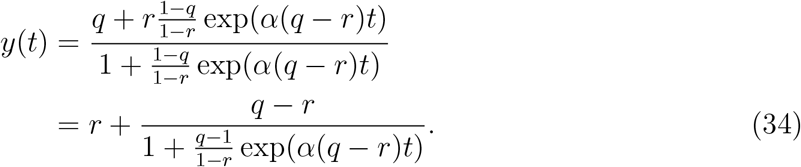

which is (6). Our next step is to write

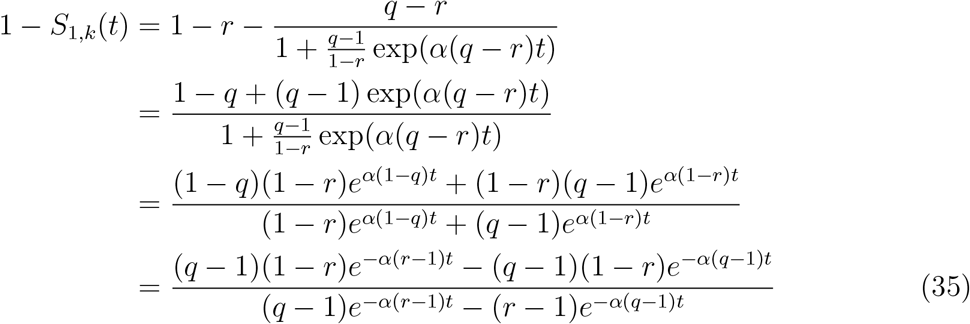

To compute *S*_2,_*_k_* (*t*) we use the recursion (7)

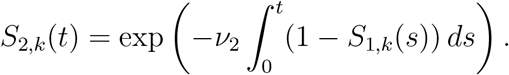

To compute integral let *f*(*t*) = (*q* − 1)*e*^−^*^α^*^(^*^r^*^−1)^*^t^* − (*r* − 1)*e*^−^*^α^*^(^*^q^*^−1)^*^t^* and note that

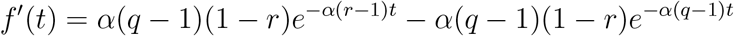

so we have

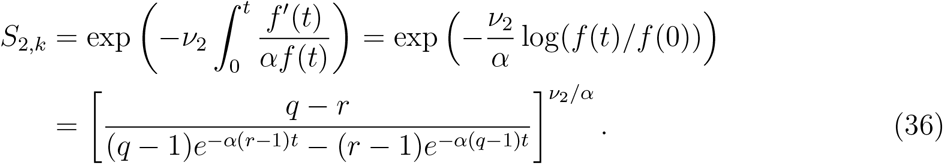

When *k* = 3, *v*_2_ = *μ*_1_ so we have

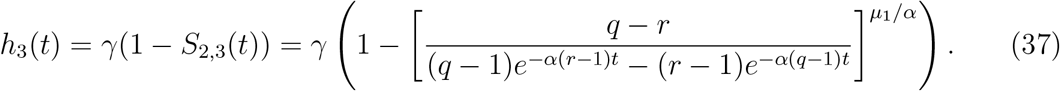

Integrating again we conclude that

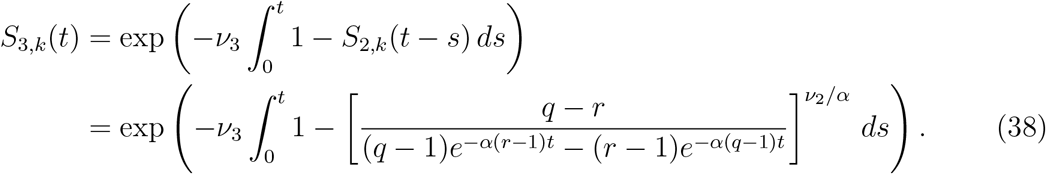

When *k =* 4, *v*_3_ = *μ*_1_ and *v*_2_ = *μ*_2_ so

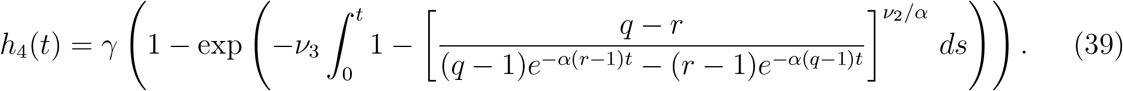

### 7.1 Approximations for *q* and *r*

When *μ* = 0 the roots *q, r* are (31)

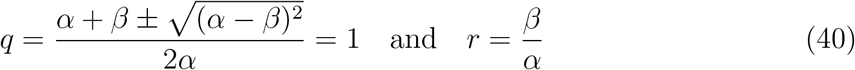

Typically the mutation rate *μ* is much smaller than *α* and *β*. When it is

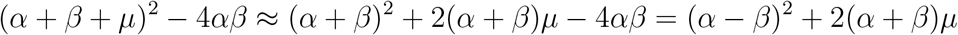

so we have

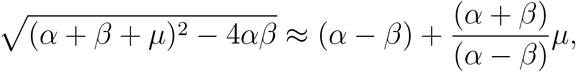

and it follows that

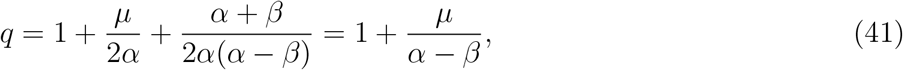

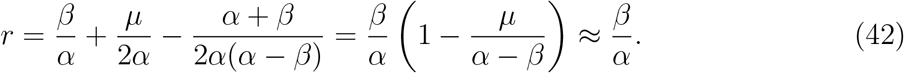

## 8 Hazard functions *H_k_*(*t*) probabilistic approach

Using (19) with the formula for *G*(*t*) given in (16)

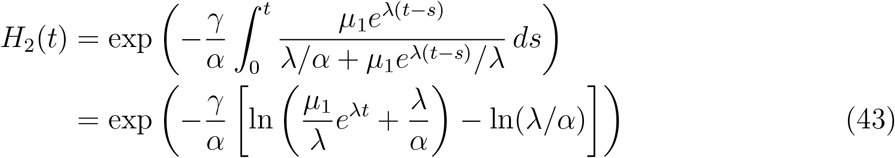

If we differentiate (43) we get

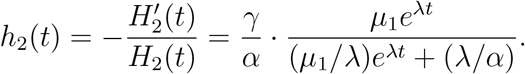

### Theorem 4.

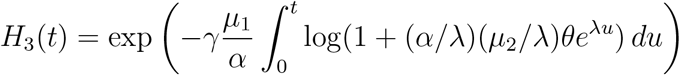

*and differentiating gives*

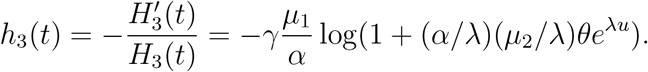

#### Proof.

Using (19) with the formula for *G*(*t*) given in (16)

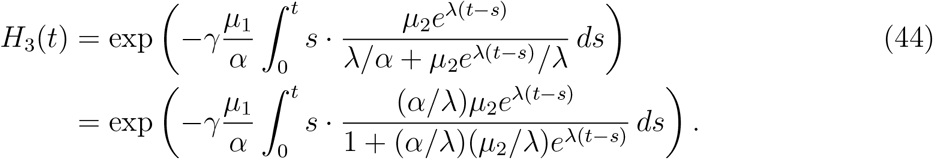

Integrating by parts with *f*(*s*) = *s* and *g*′(*s*) = the fraction under the integral

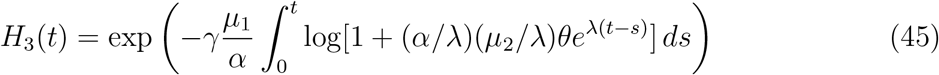

since *f*(*s*)*g*(*s*) = 0 when *s* = 0 and *s* = *t*. now change variables *u* = *t* − s.

#### Proof of

(23). If *x* is small then *x* ≈ 1 − e^−^*^x^*. Using this with

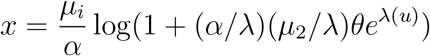

we have

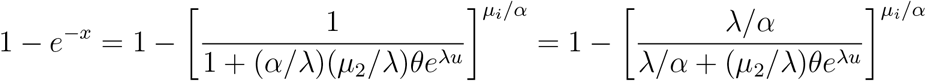

Using (18) *q* − *r* ≈ λ, and *q* − 1 ≈ *μ*_2_/λ, so the above is

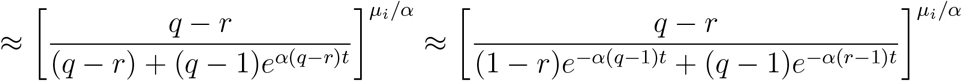

proving the desired result.

### Theorem 5.

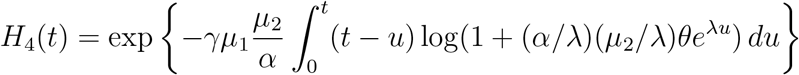

*Differentiating with respect to t*

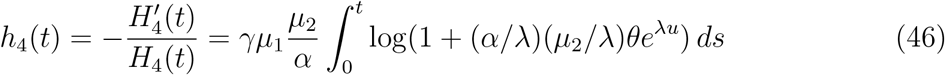

#### Proof.

Using (19) with the formula for *G*(*t*) given in (16)

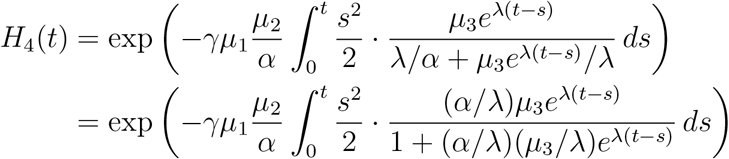

Integrating by parts with *f*(*s*) = s^2^/2 and *g*^′^(*s*) is the fraction inside the integral

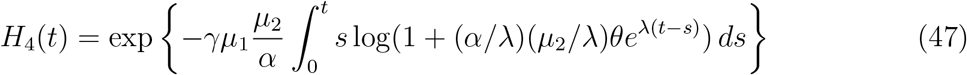

since *f*(*s*)*g*(*s*) = 0 when *s* = 0 and *s* = *t*. Changing variables *u* = *t* − *s* gives the formula for *H*_4_(*t*).

## 9 Conclusions

Here we have taken a probabilistic approach to analyze multi-stage models of cancer incidence. This leads to an intuitive proof of a simple and general formula for the distribution of the waiting time *T_k_* for the first type *k* to appear

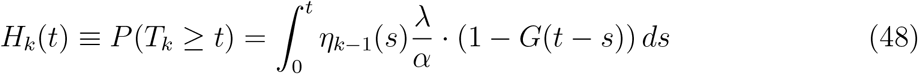

where *η_k−_*_1_(*s*) = *Nμ*_0_*μ*_1_…*μ_k−_*_1_*s^k^*^−2^/(*k* − 2)! is the rate type *k* − 1 mutations are produced at time *s*, λ/*α* is the probability a type *k* − 1 is successful, i.e., does not die out and

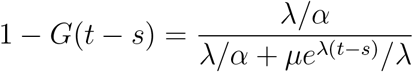

is the probability a successful type *k* − 1 born at time s produces a malignant cell by time *t*.

Differentiating (48) we can get a formula for the hazard rate 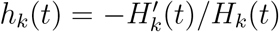. To do this it is convenient change variables *u* = *t* − *s* and write Γ*_k_*_−1_ = *Nμ*_0_*μ*_1_…*μ_k_*_−1_

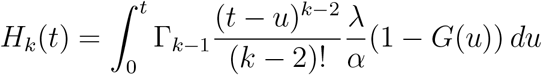

In the case *k* = 2 we have (*t* − *u*)*^k^*^−2^/(*k* − 2)! ≡ 1 so there is no *t* in the integrand and the derivative is

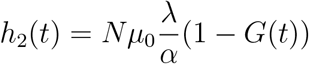

When *k* ≥ 3 we have a positive power of *t* − *u* so differentiating the upper limit does not contribute and the derivative is

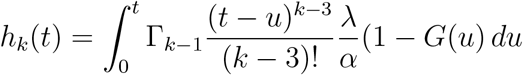

We have verified that our new formulas are almost exactly the same as the traditional ones for the *k*-stage model.

In Sections 4, 5, and 6 we considered four concrete applications that have been analyzed in the literature. In the case of pancreatic cancer, three and four stage models gave similar fits. See Figure 3. In the case of colon cancer, one gets almost identical fits from k-stage models with *k* = 2, 3, 4, 5. The parameter values for those fits (see Table 3) indicate how this is possible. The two stage fits have very small mutation rates while in the four and five stage fits, one or two mutation rates take large values. The pancreatic and colon cancer examples suggests that fitting *k*-stage models has little power to estimate the number of stages, but that power might be restored by constraining the parameter values to take on “realistic” values.

## Notes

† RD is partially supported by NSF grant DMS 1614838 from the math biology program.

